# Determining the optimal number of independent components for reproducible transcriptomic data analysis

**DOI:** 10.1101/180687

**Authors:** Ulykbek Kairov, Laura Cantini, Alessandro Greco, Askhat Molkenov, Urszula Czerwinska, Emmanuel Barillot, Andrei Zinovyev

## Abstract

**Background:** Independent Component Analysis (ICA) is a method that models gene expression data as an action of a set of statistically independent hidden factors. The output of ICA depends on a fundamental parameter: the number of components (factors) to compute. The optimal choice of this parameter, related to determining the effective data dimension, remains an open question in the application of blind source separation techniques to transcriptomic data.

**Results:** Here we address the question of optimizing the number of statistically independent components in the analysis of transcriptomic data for reproducibility of the components in multiple runs of ICA (within the same or within varying effective dimensions) and in multiple independent datasets. To this end, we introduce ranking of independent components based on their stability in multiple ICA computation runs and define a distinguished number of components (Most Stable Transcriptome Dimension, MSTD) corresponding to the point of the qualitative change of the stability profile. Based on a large body of data, we demonstrate that a sufficient number of dimensions is required for biological interpretability of the ICA decomposition and that the most stable components with ranks below MSTD have more chances to be reproduced in independent studies compared to the less stable ones. At the same time, we show that a transcriptomics dataset can be reduced to a relatively high number of dimensions without losing the interpretability of ICA, even though higher dimensions give rise to components driven by small gene sets.

**Conclusions:** We suggest a protocol of ICA application to transcriptomics data with a possibility of prioritizing components with respect to their reproducibility that strengthens the biological interpretation. Computing too few components (much less than MSTD) is not optimal for interpretability of the results. The components ranked within MSTD range have more chances to be reproduced in independent studies.

## Background

Independent Component Analysis (ICA) is a matrix factorization method for data dimension reduction [1]. ICA defines a new coordinate system in the multi-dimensional space such that the distributions of the data point projections on the new axes become as mutually independent as possible. To achieve this, the standard approach is maximizing the non-gaussianity of the data point projection distributions [1]. ICA has been widely applied for the analysis of transcriptomic data for blind separation of biological, environmental and technical factors affecting gene expression [2–6].

The interpretation of the results of any matrix factorization-based method applied to transcriptomics data is done by the analysis of the resulting pairs of metagenes and metasamples, associated to each component and represented by sets of weights for all genes and all samples, respectively [6,7]. Standard statistical tests applied to these vectors can then relate a component to a reference gene set (e.g., cell cycle genes), or to clinical annotations accompanying the transcriptomic study (e.g., tumor grade). The application of ICA to multiple expression datasets has been shown to uncover insightful knowledge about cancer biology [3,8]. In [3] a large multi-cancer ICA-based metaanalysis of transcriptomic data defined a set of metagenes associated with factors that are universal for many cancer types. Metagenes associated with cell cycle, inflammation, mitochondria function, GC-content, gender, basal-like cancer types reflected the intrinsic cancer cell properties. ICA was also able to unravel the organization of tumor microenvironment such as the presence of lymphocytes B and T, myofibroblasts, adipose tissue, smooth muscle cells and interferon signaling. This analysis shed light on the principles underlying bladder cancer molecular subtyping [3].

It has been demonstrated that ICA has advantages over the classical Principal Component Analysis (PCA) with respect to interpretability of the resulting components. The ICA components might reflect both biological factors (such as proliferation or presence of different cell types in the tumoral microenvironment) or technical factors (such as batch effects or GC-content) affecting gene expression [3,5]. However, unlike principal components, the independent components are only defined as local minima of a non-quadratic optimization function. Therefore, computing ICA from different initial approximations can result in different problem solutions. Moreover, in contrast to PCA, the components of ICA cannot be naturally ordered.

To improve these aspects, several ideas have been employed. For example, an *icasso* method has been developed to improve the stability of the independent components by: (1) applying multiple runs of ICA with different initializations; (2) clustering the resulting components; (3) defining the final result as cluster centroids; and (4) estimating the compactness of the clusters [9]. The resulting components can be then naturally ordered from the most stable to the least stable ones. This ranking is usually different from more commonly used independent component rankings based on the value of the used non-gaussianity measure (such as kurtosis) or the variance explained by the components.

The fundamental question is the determination of the number of independent components to produce. This problem can be split into two parts: a) what dimension should be selected for reducing the transcriptomic data before applying ICA (determining the effective data dimension); and b) which is the most informative number of components to use in the downstream analysis?

Determining the optimal effective data dimension for application of signal deconvolution was a subject of research in various fields. For example, ICA appeared to be a powerful method for analyzing the fMRI (functional magnetic resonance) data [9–12]. In this field, it was shown that choosing a too small effective data dimension might generate “fused components,” not reflecting the heterogeneity of the data, leading to a loss of interesting sources (under-decomposition). At the same time, choosing the effective dimension too high might lead to signal-to-noise ratio deterioration, overfitting and splitting of the meaningful components (over-decomposition) [10–12]. The influence of the effective dimension choice on the ICA performance has not been well studied in the context of transcriptomic data analysis. For example, in [3] each dataset was decomposed into a number of components in an *ad hoc* manner (n=20).

Several theoretical approaches for estimating effective data dimension exist. The simplest ones, developed for PCA analysis, are represented by the Kaiser rule aimed at keeping a certain percentage of explained variance and the broken stick model of resource distribution [13]. More sophisticated approaches employ the information theory (e.g., Akaike’s information or Minimal Description Length criteria) [13] or investigate the local-to-global data structure organization [14]. Also, computational approaches based on cross-validation have been suggested in the literature [15]. Specifically for ICA analysis, few methods have been proposed to optimize the effective dimension. For example, the Bayesian Information Criterion (BIC) can be applied to the Bayesian formulation of ICA for selecting the optimal number of components [16].

Although many of the above theoretical methods are “parameter-free,” selecting the best method for choosing an effective dimension for transcriptomic data can be challenging in the absence of a clearly defined validation strategy. One possible approach to overcome this limitation is to apply the same computational method to multiple transcriptomic datasets derived from the same tissue and disease. In this situation, it is reasonable to expect that a matrix factorization method should detect similar signals in all datasets. By taking advantage of the rich collection of public data such as The Cancer Genomic Atlas (TCGA) [17] and Gene Expression Omnibus [18], it is possible to compare and contrast the parameters of different gene expression analysis methods such as ICA.

In this study, we used TCGA pan-cancer (32 different cancer types) transcriptomic datasets and a set of six independent breast cancer transcriptomic datasets to evaluate the effect of the number of computed independent components on reproducibility and biological interpretability of the obtained results. We evaluated the reproducibility of ICA on three aspects: First, we analyzed the <u>*stability*</u> of the computed components with respect to multiple runs of ICA; second, we analyse the <u>*conservation*</u> of the computed components by varying the choice of the reduced data dimension; and third, we consider the <u>*reproducibility*</u> of the resulting set of ICA metagenes across multiple independent datasets. Our reproducibility analysis thus explores 13,027 transcriptomic profiles in 37 transcriptomic datasets, for which more than 100,000 ICA decompositions have been computed.

We finally defined a novel criterion adapted for choosing the effective data dimension for ICA analysis of gene expression, while taking into account the global properties of transcriptomic multivariate data. The Maximally Stable Transcriptome Dimension (MSTD) is defined as the maximal dimension where ICA does not yet produce a large proportion of highly unstable signals. By numerical experiments, we showed that components ranked by stability within the MSTD range tend to be more reproducible and easier to interpret than higher-order components.

## Results

### Definition of component reproducibility measures used in this study

<u>*Stability*</u> of an independent component, in terms of varying the initial starts of the ICA algorithm, is a measure of internal compactness of a cluster of matched independent components produced in multiple ICA runs *for the same dataset and with the same parameter set but with random initialization*. The exact index used for quantifying the clustering is documented in the Methods section. <u>*Conservation*</u> of an independent component in terms of choosing various orders of ICA decomposition is a correlation between matched components computed in two ICA decompositions of different orders (reduced data dimensions) *for the same dataset*. <u>*Reproducibility*</u> of an independent component is an (average) correlation between the components that can be matched after applying the ICA method using the same parameter set but *for different datasets*. For example, if a component is reproduced between the datasets of the same cancer type, then it can be considered a reliable signal less affected by technical dataset peculiarities. If the component is reproduced in datasets from many cancer types, then it can be assumed to represent a universal cancerogenesis mechanism, such as cell cycle or infiltration by immune cells. The details on computing correlations between components from different datasets are described in Methods.

### Maximally Stable Transcriptome Dimension (MSTD), a novel criterion for choosing the optimal number of ICs in transcriptomic data analysis

We used 37 transcriptomic datasets to analyze the stability and reproducibility of the ICA results conditional on the chosen number of components. ICA has been applied separately to 37 cancer transcriptomic datasets following the ICA application protocols as described in Methods.

The proposed protocol depends on a fundamental parameter *M* (effective dimension of the data and, at the same time, the number of computed independent components) whose effect on the stability of the ICs is investigated. For each transcriptomic dataset, the range of M values 2-100 has been considered. For each value of M, the data dimension is reduced to M by PCA and then data whitening is applied. Subsequently, the actual signal decomposition is applied in the whitened space by defining M new axes, where each maximizing the non-gaussianity of data point projections distribution.

For transcriptomic data, ICA decomposition provides: (a) M metagenes ranked accordingly to their stability in multiple runs (n=100) of ICA; and (b) a profile of stability of the components (set of *M* numbers in [0,1] range in descending order). Considering the largest dataset METABRIC as an example, the behavior of the stability profile as a function of M is reported in Figure 1A. The results for stability analysis for other breast cancer datasets are similar (See Supplementary Figure S2). To recapitulate the behaviour of many stability profiles, the average stability of the first k top-ranked components S_M_(k) is used (See Figure 1B). For k=M, the average stability of all computed components is denoted as S_M_^total^. Three major conclusions can be made from Figure 1. First, the average stability of the computed components S_M_^total^ decreases with the increase of M, while the average stability of the first few top ranked components, i.e., S_M_(10), weakly depends on M (Figure 1B). Moreover, S_M_^total^ is characterized by the presence of local maxima, defining certain distinguished values of M that correspond to the (locally) maximally stable set of components (Figure 1B). Third, the stability profiles for various values of M can be classified into those for which the stability values are distributed approximately uniformly and those (usually, in higher dimensions) forming a large proportion of the components with low stability (*I*_*q*_ between 0.2 and 0.4) (Figure 1A).

**Figure 1.**
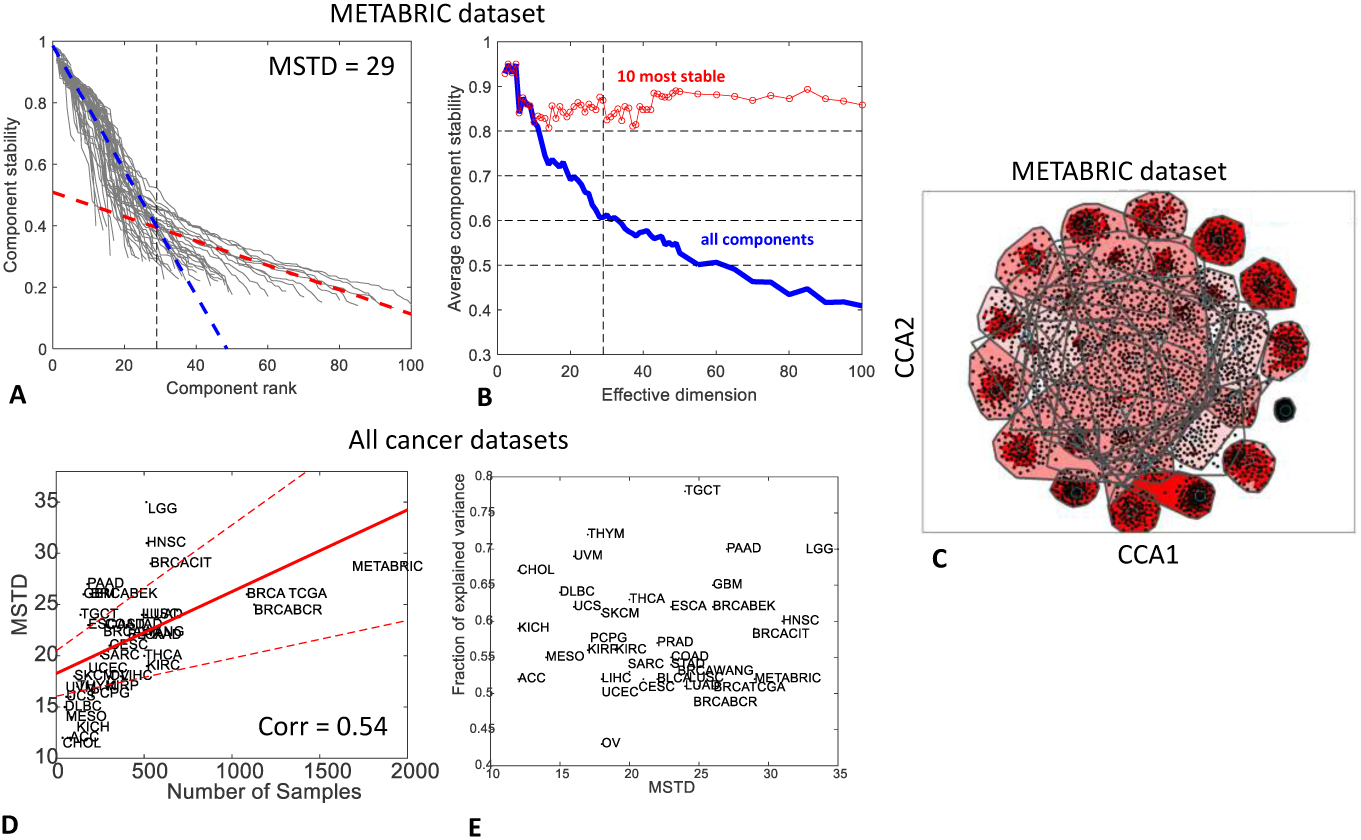
Defining Maximally Stable Transcriptomic Dimension (MSTD) value in 37 transcriptomics cancer datasets (13027 samples in total). In A-C) an example of the analysis is presented for the largest breast cancer dataset METABRIC. A) stability profiles for ICA decompositions in various dimensions (from 2 to 100) shown by grey lines. Two-line clustering result is shown by blue and red dashed lines, with MSTD determined as the point of their intersection (vertical dashed line). B) average stability profile S_M_^total^ (blue line) and the average stability of 10 most stable components S_M_(10) (red line). C) visualizing the results of computing ICA 100 times with MSTD=29 components in the METABRIC dataset and component clustering (*icasso* package, Canonical Correlation Analysis (CCA) plot). Each black point represents a component, red lines show significant correlations between them, polygons show the convex hull area of the clusters. D) Dependence of MSTD on the number of samples for all datasets. E) Dependence of the fraction of explained variance on MSTD for all datasets.

Considering these observations, we hypothesized that the optimal number of independent components -- large enough to avoid fusing meaningful components and yet small enough to avoid producing an excessive amount of highly unstable components -- should correspond to the inflection point in the distribution of the stability profiles (Figure 1A). To find this point, the stability measures have been clustered along two lines, which is analogous of 2-means clustering but with lines as centroids. In this clustering, the line with a steeper slope (Figure 1A, blue line) grouped the stability profiles with uniform distribution, while another line (Figure 1A, red line) matched the mode of low stability components. The intersection of these lines provided a consistent estimate of the effective number of independent components. We call this estimate Maximally Stable Transcriptome Dimension (MSTD) and in the following we investigated its properties. We note that, as in various information theory-based criteria (BIC, AIC), this estimate is free of parameters (thresholds), and it only exploits the property of the qualitative change in the character of the stability profile in higher data dimensions for transcriptomic data.

In most of the cancer transriptomics datasets used in our analysis, MSTD was found to correspond roughly to the average stability profile S_M_^total^ ≈ 0.6 (Supplementary Figure S2). In Figure 1D, the dependence of MSTD on the number of samples contained in the transcriptomic dataset is investigated for all the 37 transcriptomic datasets. As shown in Supplement Figure 1, MSTD increased with the number of samples; however, this trend was weaker than other estimates of an effective dimension such as Kaiser rule and broken stick distribution-based data dimension estimates. Finally, the fraction of variance explained by the linear subspace spanned by MSTD number of components was evaluated (Figure 1E), and it was observed that the fraction of variance explained varied from 0.45 to 0.75 with a median of 0.56.

### Underestimating the effective dimension (M<MSTD) leads to a poor detection of known biological signals

Previous large-scale ICA-based meta-analyses [3] have shown that some of the ICs derived from the decomposition of a cancer transcriptomic data were clearly and uniquely associated with known biological signals. For example, one of these signals was the one connected to proliferative status of tumors. Another example was given by the signals related to the infiltration of immune cells that were also strongly heterogeneous across cancer patients.

We have checked the reproducibility of several metagenes results from previous meta-analyses [3] where all ICA decompositions were treated as a function of M. For this analysis, we employed the METABRIC breast cancer dataset, which was not included in the input data of the previous publication [3] and thus it had not been used to derive the metagenes of that work. In addition, we checked how the significance of intersections between the genes defining the components and several reference gene sets (produced independently of the ICA analyses) behaved as a function of M.

We applied the previously developed correlation-based approach to match previously identified metagenes with the ones computed for a new METABRIC dataset (see Methods section). The components were oriented accordingly to the direction of the heaviest tail of the projection distribution. When matching an oriented component to the previously defined set of metagenes, we verified that the resulting maximal correlation should be positive, i.e. large positive weights in one metagene should correspond to large positive weights in another metagene.

One of the most important case studies is reproducibility of the “proliferative” metagene in different data dimensions. It is investigated in Figure 2A-C. For this metagene, we computed correlations with M newly identified independent components. As an example, the profile of correlations for M=100 is shown in Figure 2B. It can be seen that one of the components (ranked #7 by stability analysis) is much better correlated to the proliferative metagene than any other component. Therefore, component #7 is called “best matched” in this case, for M=100, and “well separable.” Repeating this analysis for all M and reporting the observed maximal correlation coefficient and the corresponding stability value gives a plot shown in Figure 2A. Separability of the best matched component from the other components is visualized in Figure 2C.

**Figure 2.**
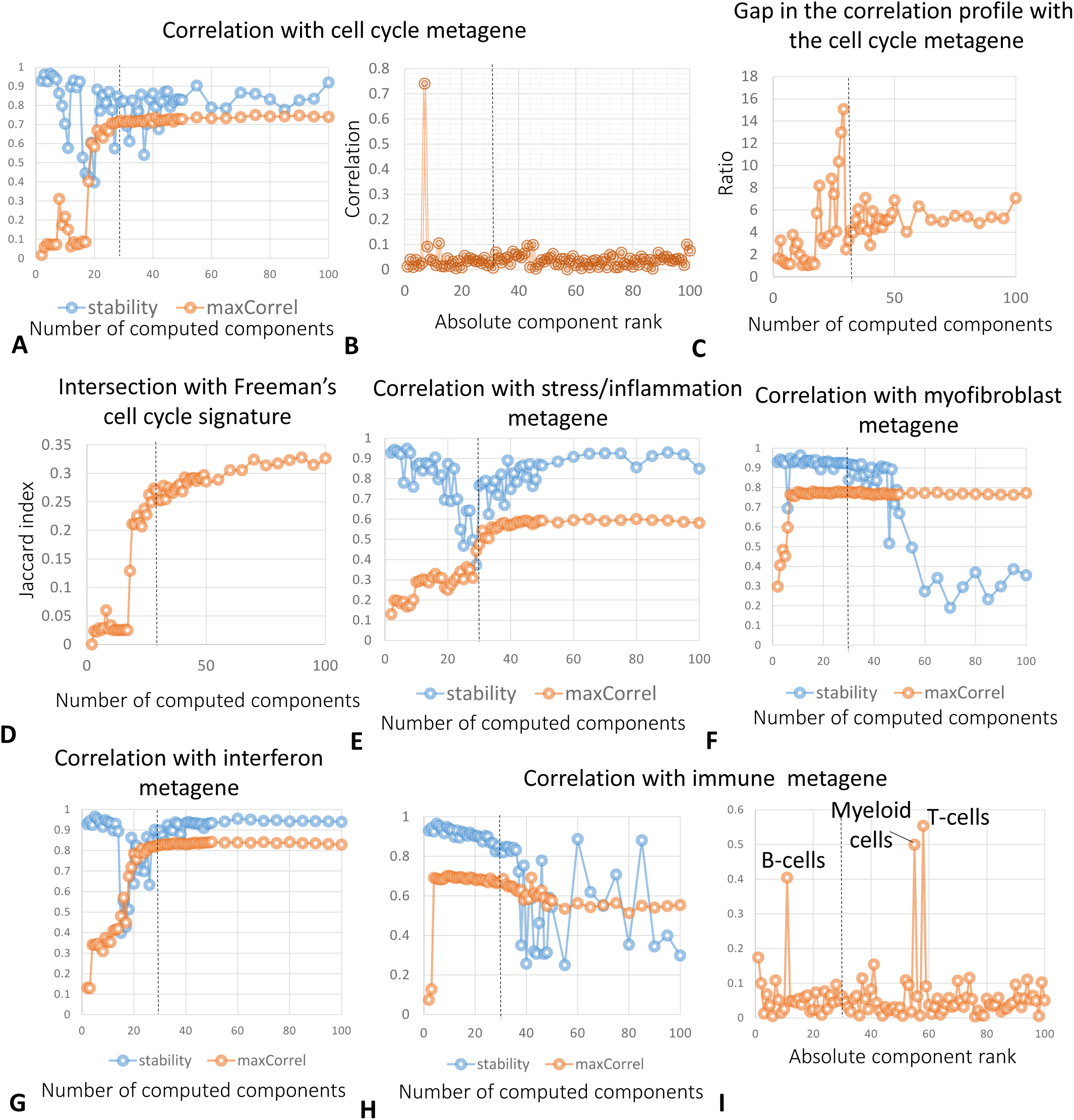
Analysis of reproducibility of previously identified metagenes in independent components of the METABRIC dataset. A,E,F,G,H) show correlations (see Methods section) with the cell cycle, inflammation, myofibroblast, interferon signaling, immune-related metagenes from [3] as a function of the chosen data dimension M, and the stability of the best matched component. C) shows the ratio between the correlation value of the proliferation metagene with the best matched component and the second best correlation (gap). D) shows an intersection (Jaccard index) of the Freeman’s cell cycle signature [19] and the set of top-contributing genes (projection>5.0) from the proliferation-associated independent component. B,I) correlation of the cell cycle and immune-related metagene with the best matched component in the M=100 ICA decomposition as a function of the stability-based component rank. In all plots, the vertical dashed line shows the MSTD value for the METABRIC dataset.

As it can be seen from Figure 1A, the biologically expected signals (i.e., cell cycle) can be poorly detected for M<MSTD; however, once the best matching component with significant correlation was found, it remained unique and was detected robustly even for very large values of M>>MSTD. For example, even when 100 components (M) were computed, the correlation between the previously defined proliferative metagene and the best matched independent component did not diminish (Figure 2A). Moreover, the separability of the best matched component from the rest of the components was not ruined (Figure 2C). In this example, the identification of cell cycle component remained clear (large and well-separated correlation coefficient) for M>>MSTD. This result was consistent and complementary when compared with the previously observed weak dependence of S_M_(10) on M. Indeed, the “proliferative” best matched component had stability rank k in the range [6,11]. That is, it remained stable in ICA decompositions in all dimensions. Moreover, the intersection of a recently established proliferation gene signature [19] with the set of top contributing genes of the best matched component improved with M and saturates (Figure 2D). This proves that the detection of the proliferation-associated signal with ICA does not depend on the ICA-based definition of the proliferative metagene.

Together with the proliferative signal, other metagenes from the previously cited ICA-based meta-analysis [3] were robustly identified in our analysis. In Figure 2 E-H, we showed the correlation with the best matching component for the metagenes associated with the presence of myofibroblasts, inflammation, interferon signaling and immune system, as a function of M. These plots illustrated different scenarios that can result from such analysis. The myofibroblast-associated metagene was robustly detected for all values of M>7 (Figure 2F). However, the stability of the best matching component was deteriorated in higher-order ICA decompositions (M>45). For the inflammation-associated metagene, an ICA decomposition with M>38 was needed to robustly detect a component that correlates with the metagene (Figure 2E).

Interestingly, the immune-associated metagene was found robustly matched starting from M=4. However, in higher-order decompositions (starting from M=30) it could be matched to several components that can be associated with specific immune system-related signals (Figure 2 H-I). Hypergeometric tests applied to the sets of top-contributing genes (weights larger than 5.0) allowed us to reliably interpret these components as being associated with the presence of three types of immune-related cells: T cells (corrected enrichment p-value=10^-39^ with “alpha beta T cells” signature [20], other immune signatures are much less significant), B cells (p-value=10^-7^ with “B cells, preB.FrD.BM” signature) and myeloid cells (p-value=10^-78^ with “Myeloid Cells, DC.11cloSer.Salm3.SI” signature).

### Overestimating the number of components (M>>MSTD) produces multiple ICs driven by small gene sets

We observed that the higher-order ICA decompositions (M>>MSTD) produced a larger number of components driven by small gene sets (frequently, one gene), such that the projections of the genes in this “outlier” set is separated by a relatively large gap with the rest of the projections. We thus designed a simple algorithm to distinguish such components driven by a small gene set from all the others. The names of the genes composing these small sets were used for annotating the corresponding components (Figure 3A, right part).

**Figure 3.**
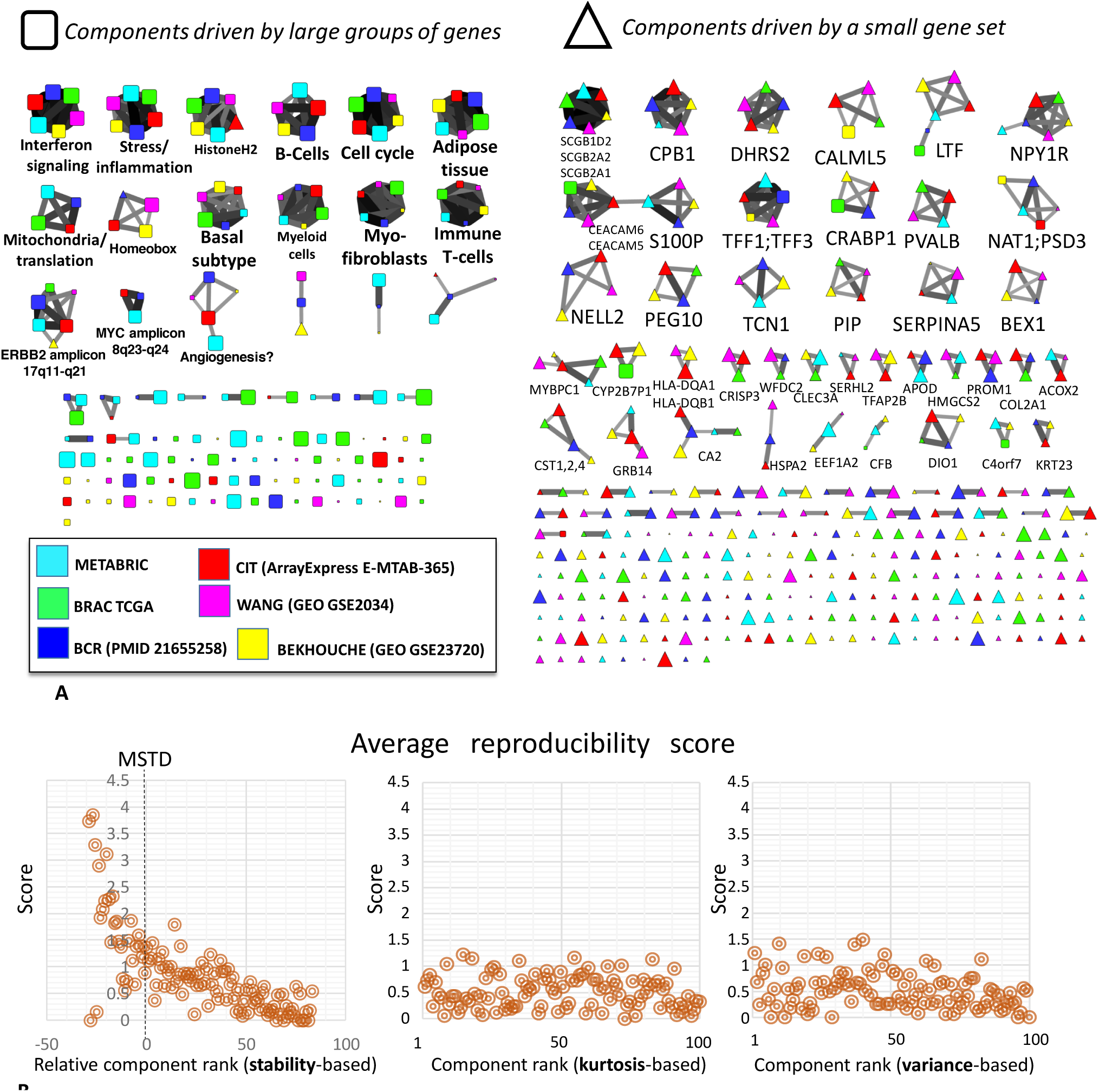
Analysis of component reproducibility in independent datasets. A) Graph of reciprocal correlations showing the reproducibility of the metagenes of independent components in 6 independent breast cancer datasets. Each node here is an independent component, represented by a metagene, from an ICA decomposition with M=100 components. Edges show only reciprocal correlations between metagenes with Pearson correlation > 0.3. Triangles (on the right) show the components driven by the expression of a small group of genes (frequently, one gene). Node size reflects the rank of the component based on the stability in multiple runs of fastICA (larger nodes are more stable ones). The edge width and the color reflect the value of the correlation coefficient between two metagenes, with thicker edges showing larger correlation values. Several pseudo-cliques of highly reproducible components are annotated either by the dominating small group of genes (pseudo-cliques of triangle nodes), or by comparing to the results of the previously published large-scale ICA-based analysis of gene expression [3] or by performing the hypergeometric test using the set of top-contributing genes (with projection larger than 5.0 onto the component). The analogous correlation graph computed for MSTD number of components is provided in Supplementary Figure SF3. B) average reproducibility score (sum of reciprocal correlation coefficients of an independent component) for the correlation graph shown in A), as a function of the relative (component rank minus MSTD value for a given dataset, for stability-based ranking) or absolute (for other ranking types) component rank. It is clear that only stability-based ranking matches the reproducibility score.

It was observed that the presence of such “small gene set-driven” components is a characteristic of higher-order ICA decompositions (M>>MSTD), much less present in ICA decompositions with M≤MSTD (compare Figure 3A and Supplementary Figure SF2).

To check the biological significance of the outlier genes, we considered as a case study the higher-order (M=100) ICA decomposition of the METABRIC breast cancer dataset. We collected all those genes found to be drivers of at least one “small gene set-driven” component. We obtained in this way a set of 98 genes listed in Supplementary Table ST2. This list appeared to be strongly enriched (p-value = 10^-12^ after correction for multiple testing) in the genes of the signature DOANE_BREAST_CANCER_ESR1_UP “Genes up-regulated in breast cancer samples positive for ESR1 compared to the ESR1 negative tumors” from Molecular Signature Database [21] and several other specific to breast cancer gene signatures. This analysis thus suggested that at least some of the identified “small gene set-driven” components are not the artifacts of the ICA decomposition, but they can be biologically meaningful and reproducible in independent datasets (Figure 3A, right part).

### Most stable components with stability rank≤MSTD have more chances to be reproduced across independent datasets for the same cancer type

It would be reasonable to expect that the main biological signals characteristic for a given cancer type should be the same when one studies molecular profiles of different independent cohorts of patients. Therefore, we expect that for multiple datasets related to the same cancer type, ICA decompositions should be somewhat similar; hence, reciprocally matching each other. We called this expected behavior “reproducibility,” and here we studied this by applying ICA to six relatively large breast cancer transcriptomic datasets. Of note, these datasets were produced using various technologies of transcriptomic profiling (Supplementary Table ST1).

To identify the reproducible components, we applied the same methodology as in the previously published ICA-based gene expression meta-analysis [3]. We decomposed the six datasets separately and then constructed a graph of reciprocal correlations between the obtained metagenes. Correlation between two sets of components is called reciprocal when a component from one set is the best match (maximally correlated) to a component from another set, and vice versa (see Methods for a strict definition).

Pseudo-cliques in this graph, consisting of several nodes, correspond to reproducible signals detected by ICA. As shown in Figure 3, multiple reproducible signals were identified in the analysis. Some of them correspond to signals already identified in [3] (e.g., cell cycle, interferon signaling, microenvironment-related signals), and some correspond to newly discovered biological signals (e.g., ERBB2 amplicon-associated). Some other pseudo-cliques are associated with “small gene set-driven” components (frequently, one gene-driven), such as TFF1-3-associated or SCGB2A1-2-associated components.

The genes driver of reproducible and “small gene set-driven” components (S100P, TFF1, TFF3, SCGB2A1, SCGB1D2, SCGB2A2, LTF, CEACAM6, CEACAM5 being most remarkable examples) have been investigated in detail, to further check their biological interest. They were found to be the genes known to be associated with breast cancer progression [22]. For example, seven of the nine previously mentioned genes form a part of a gene set known to be up-regulated in the bone relapses of breast cancer (M3238 gene set from MSigDB).

To quantify the reproducibility of the components, we computed a reproducibility score. It is a sum of correlation coefficients between the component and all reciprocally correlated components from other datasets. By construction, the maximum value of the score is 5, which meant that a component with such a score would be perfectly correlated with the reciprocally related components from five other datasets. We studied the dependence of this score as a function of the relative to MSTD component stability-based rank (Figure 3B). From this study, it follows that even for the high-order ICA decompositions, the components ranked by their stability within MSTD range, have an increased likelihood of being reproduced in independent datasets collected for the same cancer type.

To show that the stability-based ranking of genes is more informative compared with the standard rankings of independent components, we performed a computational analysis in which we compared the stability-based ranking with the rankings based on non-gaussianity (kurtosis) and explained variance. These two measures are frequently used to rank the independent components [6]. From Figure 3B it is clear that the stability-based ranking of independent components corresponds well to the reproducibility score, while two other simpler measures do not.

It can also be shown that the total number of reciprocal correlations with relatively large correlation coefficients (|r|>0.3) between ICA-based metagenes computed for several independent datasets is significantly bigger when the component stabilization approach is applied (Supplementary Figure S4). This proves the utility of the applied stabilization-based protocol of ICA application to transcriptomic data.

### Computing large number of components (M>>MSTD) does not strongly affect the most stable ones

We lastly used ICA decompositions of 37 transcriptomic datasets to compare the ICA decompositions corresponding to M=MSTD with the higher-order decompositions, M=50 or M=100.

It was found that the components calculated in lower data dimensions can be relatively well matched to the components from higher-order ICA decompositions (Figure 4). More precisely, 90% of the components defined for M=MSTD had a reciprocal best matched component in the M=100 ICA decomposition. Most stable components had a clear tendency to be reproduced with high correlation coefficient (r>0.8). Only 10% of the components had only non-reciprocal or too small correlations between two decompositions (in other words, *not conserved* in higher-order ICA decompositions).

**Figure 4.**
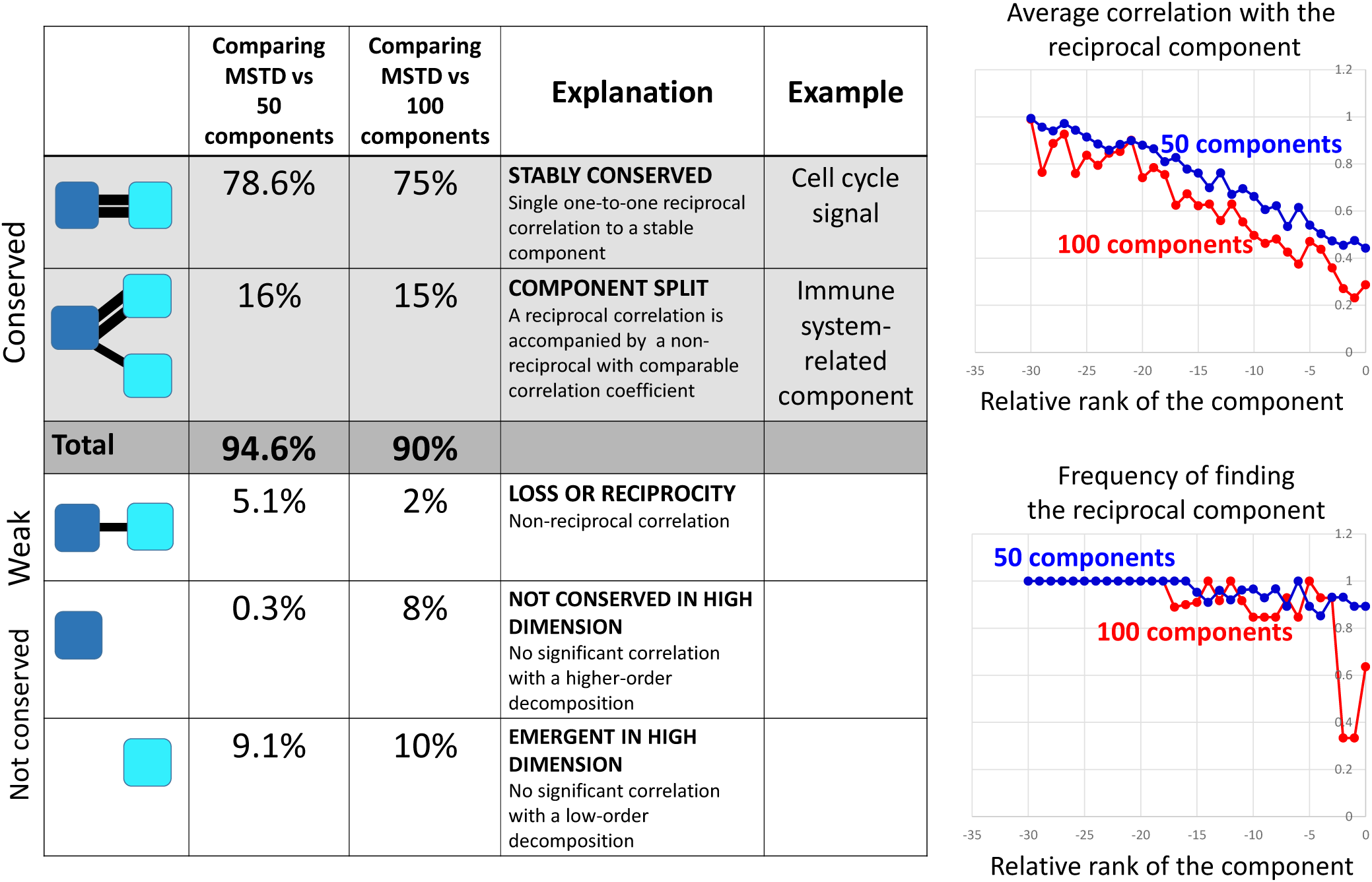
Conservation or non-conservation of independent components in higher than MSTD dimensions. ICA decomposition in MSTD data dimension is compared with the one of dimension 50 and 100. Table on the left: list of different scenario with the relative frequency of each of them estimated for a pan-cancer TCGA dataset. Plots on the right show the frequency of finding the reciprocally correlated component in the higher dimension and the average correlation coefficient of the reciprocal correlation, as a function of the relative rank of the component (component rank minus MSTD value for a given dataset).

Approximately 15% of the components in M=MSTD ICA decomposition together with reciprocal maximal correlation also had a non-reciprocal correlation to one of the components in M=100 ICA decomposition (Figure 4). This case can be described as splitting a component into two or more components in the higher-order ICA decompositions. At least one such split had a clear biological meaning, namely the splitting of the component representing the generic “immune infiltrate.” The resulting “split” components more specifically represented the role of T cells, B cells and myeloid cells in the tumoral microenvironment (see the “*Underestimating the effective dimension…*” Results section).

## Discussion

Our results shed light on the organization of the multivariate distribution of gene expression in the high-dimensional space. It appears that the organization contained two relatively well separated parts: *the dense one* of a relatively small effective dimension and *the sparse one*. The former contained the genes from within co-regulated modules that contained from few tens to few hundreds of genes. The latter was spanned by the genes with unique regulatory programs (perhaps tissue-specific) weakly shared by the other genes. Here the sparsity was understood in the sense of low local multivariate distribution density.

Independent Component Analysis can capture both these parts of the multivariate distribution. However, while the dense part defined independent components with approximately uniformly distributed stabilities, starting from highly stable to less stable, the sparse part was spanned by the components characterized mostly by small stability values.

This organization of the gene expression space is captured in the distribution of ICA stability profiles for varying M, which allowed us to define the Maximally Stable Transcriptome Dimension (MSTD) value, roughly reflecting the dimension of the dense part of the gene expression distribution. In one hand, when underdecomposing (compressing too much by dimension reduction, M<MSTD) a transcriptomic dataset, the resulting independent components are hard to interpret. In the other hand, overdecomposing transcriptomes (choosing the effective dimension much bigger than MSTD) is not dramatically detrimental: one can choose to explore a relatively multi-dimensional subspace of a transcriptomic dataset, taking into account that applying matrix factorization methods in higher dimensions becomes computationally challenging and prone to bad algorithm convergence. Nevertheless, higher-order decompositions might allow capturing the behavior of some tissue-specific or cancer type-specific biomarker genes from the sparse part of the distribution, which can be found reproducible in other independent studies.

In our computational experiments, we selected 100 as the maximum order of ICA decomposition (M) to test. However it is possible to examine even higher orders of ICA decompositions, reducing the data to more than 100 dimensions, but not more than the total number of samples, of course. In practice, computing ICA in such high dimension leads to significant deterioration of the fastICA algorithm convergence, so exploring M>100 might be too expensive in terms of computational time. Moreover, our study suggests that the most interesting for interpretation components are usually positioned within the first few ten top ranks: therefore, 100 seems to be a reasonable limit for dimension reduction when applying ICA to transcriptomic data.

Our proposed approach can be used for comparing intrinsic reproducibility, at different levels, of various matrix factorization methods. For example, it would be of interest to compare the widely used Non-negative matrix factorization (NMF) method [6,7] with ICA to assess reproducibility of extracted metagenes in independent datasets of the same nature.

More generally, systematic reproducibility analysis can be a useful approach for establishing the best practices of application of the bioinformatics methods.

## Conclusion

By using a large body of data and comparing 0.1 million decompositions of transcriptomic datasets into the sets of independent components, we have checked systematically the resulting metagenes for their reproducibility in several runs of ICA computation (measuring *stability*), for their reproducibility between a lower order and higher-order ICA decompositions (*conservation*), and between metagene sets computed for several independent datasets, profiling tumoral samples of the same cancer type (*reproducibility*).

From the first of such analyses, we formulated a minimally advised number of dimensions to which a transcriptomic dataset should be reduced called Maximally Stable Transcriptome Dimension (MSTD). Reducing a transcriptomic dataset to a dimension below MSTD is not optimal in terms of the interpretability of the resulting ICA components. We showed that for relatively large transcriptomic datasets, MSTD could vary from 15 to 30 and that the number of samples matters relatively weakly.

From the second analysis, we concluded that the suggested protocol of ICA application to transcriptomic data is conservative, i.e., the components identified in a higher dimension (for example, in one hundred dimensional space) can be robustly matched with those components obtained in the dimensions comparable with MSTD. Moreover, we described an effect of interpretable component splitting in higher dimensions, leading to detection of finer-grained signals (e.g., related to the decomposition of the immune infiltrate in the tumor microenvironment). At the same time, the application of ICA in high dimensions resulted in a greater proportion of unstable components, many of them were driven by expression of small (one to three members) gene sets. Yet, some of these small gene set-driven components were highly reproducible and biologically meaningful.

From the third analysis, we established that the used protocol of ICA application, with ranking the independent components based on their stability, prioritized those components having more chances to be reproduced in independent transcriptomic datasets. Moreover, when ICA was applied in higher dimensions, the components within the MSTD range still have more chances to be reproduced.

In sum, our results confirmed advantageous features of ICA applied to gene expression data from different platforms, leading to interpretable and quantifiably reproducible results. Comparing ICA analyses performed in various dimensions and multiple independent datasets for the same cancer types allow prioritizing of the most reliable and reproducible components which can be quantitatively recapitulated in the form of metagenes or the sets of top contributing genes. We expect that ICA will demonstrate similar properties in other large-scale transcriptomic data collections such as scRNA-seq data.

## Materials and Methods

### Transcriptomics cancer data used in the analysis

Expression data derived for 32 solid cancer types (ACC, BLCA, BRCA, CESC, CHOL, COAD, DLBC, ESCA, GBM, HNSC, KICH, KIRC, KIRP, LGG, LIHC, LUAD, LUSC, MESO, OV, PAAD, PCPG, PRAD, READ, SARC, SKCM, STAD, TGCT, THCA, THYM, UCEC, UCS, UVM) were downloaded from the TCGA web-site and internally normalized. Normalized breast cancer datasets from CIT, BCR, WANG, BEKHOUCHE were re-used from the previous study [3]. Normalized METABRIC breast cancer expression dataset was downloaded from cBioPortal at this link http://www.cbioportal.org/study?id=brca_metabric. When it was not already the case, the data values were converted into logarithmic scale.

The list of breast cancer transcriptomic datasets used for reproducibility study is available in Supplementary Table ST1.

### ICA decompositions computation

We applied the same protocol of application of ICA decomposition as in [3]. In the ICA decomposition X ≈ AS, X is the gene expression (sample vs gene) matrix, A is the (sample vs. component) matrix describing the loadings of the independent components, and S is the (component vs. gene matrix) describing the weights (projections) of the genes in the components. To compute ICA, we used the *fastICA* algorithm [1] accompanied by the *icasso* package [23] to improve the components estimation and to rank the components based on their stability. ICA was applied to each transcriptomic dataset separately.

For each analysed transcriptomic dataset, we computed *M* independent components (ICs), using *pow3* nonlinearity and *symmetrical* approach to the decomposition, where M = [2…50, 55, 60, 65, 70, 75, 80, 85, 90, 95, 100]. In those cases, when M exceeded the total number of samples, the maximum M was chosen equal to 0.9 multiplied by the number of samples (moderate dimension reduction improves convergence). We found that the MATLAB implementations of *fastICA* performs superior to other implementations (such as those provided in *R* [25]). The computational time required for performing all the 0.1 million ICA decompositions used in this study is estimated in ∼1500 single processor hours using MATLAB while other implementations would not make this analysis feasible at all. In our analysis, we used Docker with packaged compiled MATLAB code for fastICA together with MATLAB Runtime environment, which can be readily used in other applications and does not require MATLAB installed [26]. An example of computational time needed for the analysis of two transcriptomic datasets of typical size (full transcriptome, from 200 to 1000 samples) is provided in Supplementary Figure SF5. As a rough estimate, it takes 3 hours to analyze a transcriptomic dataset with 200 samples and 7 hours to analyze a dataset with 1000 samples, using an ordinary laptop. In each such analysis, more than 2000 ICA decompositions of different orders have been made.

### The algorithm for determining the Most Stable Transcriptome Dimension (MSTD)

1. Define two numbers [*Mmin*, *Mmax*] as the minimal and maximal possible numbers of the computed components.
2. Define the number *K* of ICA runs for estimating the components stability. In all our examples, we used K=100.
3. For each *M* between *M*_*min*_ and *M*_*max*_ (or, with some step) do
  3.1) Compute *K* times the decomposition of the studied dataset into *M* independent components using the *fastICA* algorithm. This results in computation of *M*x*K* components.
  3.2) Cluster *M*x*K* components into *M* clusters using agglomerative hierarchical clustering algorithm with the measure of dissimilarity equal to 1-|*r*_*ij*_|, where *r*_*ij*_ is the Pearson correlation coefficient computed between components.
  3.3) For each cluster *C*_*k*_ out of *M* clusters (*C*_*1*_*, C*_*2*_*,…, C*_*N*_) compute the stability index using the following formula

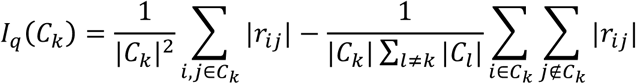

where |*C*_*k*_| denotes the size of the *k*th cluster.
  3.4) Compute the average stability index for *M* clusters:

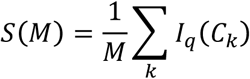
4. Select the MSTD as the point of intersection of the two lines approximating the distribution of stability profiles (Figure 1A). The lines are computed using a simple k-lines clustering algorithm [27] for k=2, implemented by the authors in MATLAB, with the initial approximations of the lines matching the abscissa and the ordinate axes of the plot.

The index used in 3.3 is a widely used index of clustering quality defined as a difference between the average intra-cluster similarity and the average inter-cluster similarity. In [9] this index was introduced to estimate the quality of clustering of independent components after multiple runs with random initial conditions, and tested in application to fMRI data. In the case of clustering independent components, *I*_*q*_ = 1 corresponds to the case of perfect clustering of components such that all the components in one cluster are correlated with each other with |*r*| = 1, and that all components in the same cluster are orthogonal to any other component (in the reduced and whitened space).

### Comparing metagenes computed for different datasets and in different analyses

Following the methodology developed previously in [3], the metagenes computed in two independent datasets were compared by computing a Pearson correlation coefficient between their corresponding gene weights. Since each dataset can contain a different set of genes, the correlation is computed on the genes which are common for a pair of datasets. Note that this common set of genes can be different for different pairs of datasets. The same correlation-based comparison was done with previously defined and annotated metagenes. We computed the correlation only between those genes having projection value more than 3 standard deviations in the identified component.

When comparing two sets of metagenes **A** = {A_1_,…,A_M_} and **B** = {B_1_,….,B_N_}, in order to do component matching, we focused on the maximal correlation of a metagene from one set with all components from another set. If B_i_ = arg max(corr(A_j_,**B**)) then Bi is called *best matched*, for Aj, metagene from the set **B**. If B_i_ = arg max(corr(A_j_,**B**)) and A_j_ = arg max(corr(B_i_,**A**)), then the correlation between B_i_ and A_j_ is called *reciprocal*.

In all correlation-based comparisons, the absolute value of the correlation coefficient was used.

The orientation of independent components was chosen such that the longest tail of the data projection distribution would be on the positive side. Then, for quantifying an intersection between a metagene and a reference set of genes (e.g., cell cycle genes), simple Jaccard index was computed between the reference gene set and the set of top-contributing genes to the component, with positive weights >5.0.

### Determining if a small gene set is driving an independent component

To distinguish whether an independent component is driven by a small gene set, the distribution of gene weights *W*_*i*_ from the component was analyzed. For each tail of the distribution (positive and negative), the tail weight was determined as the total absolute sum of weights of the genes exceeding certain threshold W^top^. The heaviest tail of the distribution was identified as the tail with the maximum weight. For the heaviest tail and for the set of genes *P* with absolute weights exceeding W^top^, sorted in descending order by absolute value, we studied the gap distribution of values G_i_ = W_i_/W_i+1_, i1 P. If there was a single value of G_i_ exceeding a threshold G^max^, then the component was classified as being driven by a small set of genes corresponding to the indices {i; i ≤ max(k; G_k_ ≤ G^max^)}. The values W^top^ = 3.0, G^max^ = 1.5 collected the maximal gene set size = 3 in all ICA decompositions. These are few genes with atypically high weights separated by a significant gap from the rest of the distribution (note that these genes cannot always be considered outliers since they and the resulting independent components can be reproducible in independent datasets).

## Declarations

### List of abbreviations

ICA: Independent Component Analysis
IC: Independent Component

### Ethics approval and consent to participate

Not applicable

### Consent for publication

Not applicable

### Availability of data and materials

This study is based on the analysis of public data. The provenance of the public data used in this study is indicated in the Method section and Supplementary Table ST1.

## Acknowledgements

We thank Dr. Anne Biton for sharing the normalized public transcriptomics data for four breast cancer datasets. We also thank Prof. Joseph H. Lee (Columbia University) for critical reading and improving the manuscript text.

## Authors’ contribution

UK LC EB AZ designed the study and developed the methodology, UK LC AG AM UC AZ performed the computational experiments, UK LC UC AZ wrote the manuscript, all authors read and edited the manuscript.

## Supplementary Figures

**Supplementary Figure SF1.**
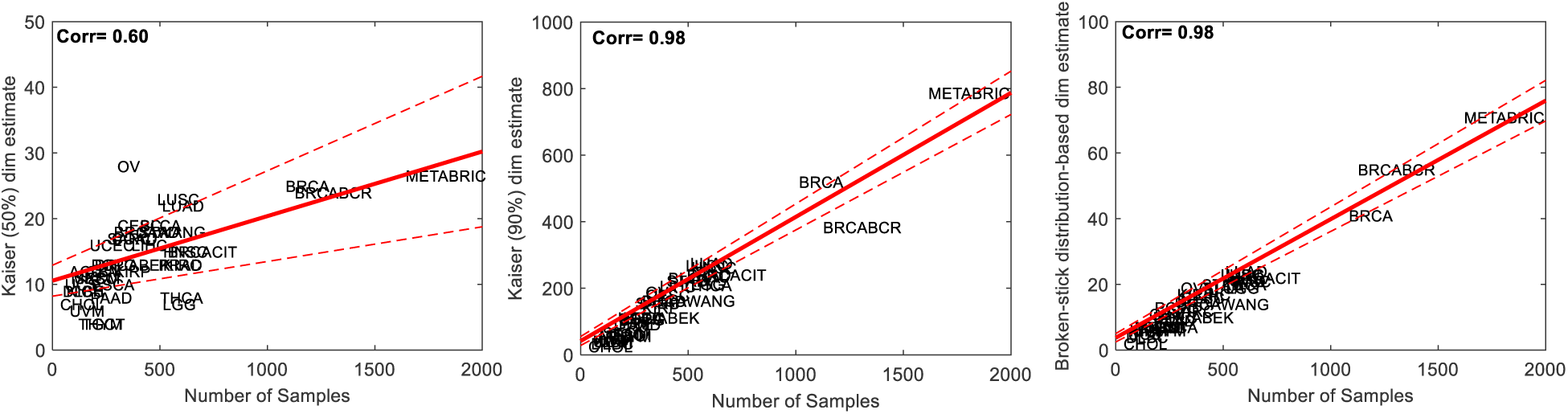
Standard estimations of intrinsic dimensionality (by Keiser rule or by broken stick distribution) of cancer datasets.

**Supplementary Figure SF2.**
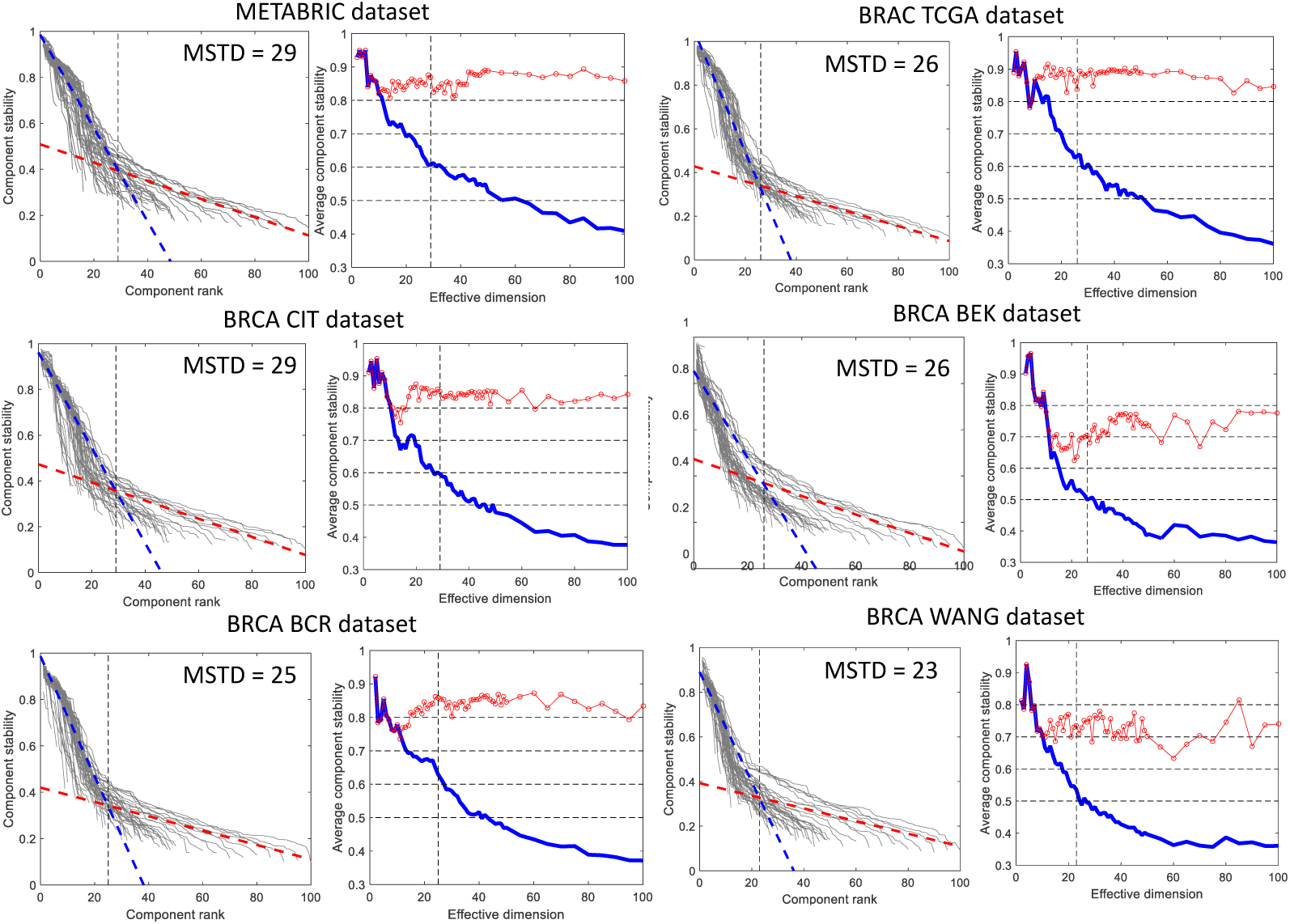
Estimating MSTD dimension for six breast cancer datasets. The notations are the same as in Figure 1.

**Supplementary Figure SF3.**
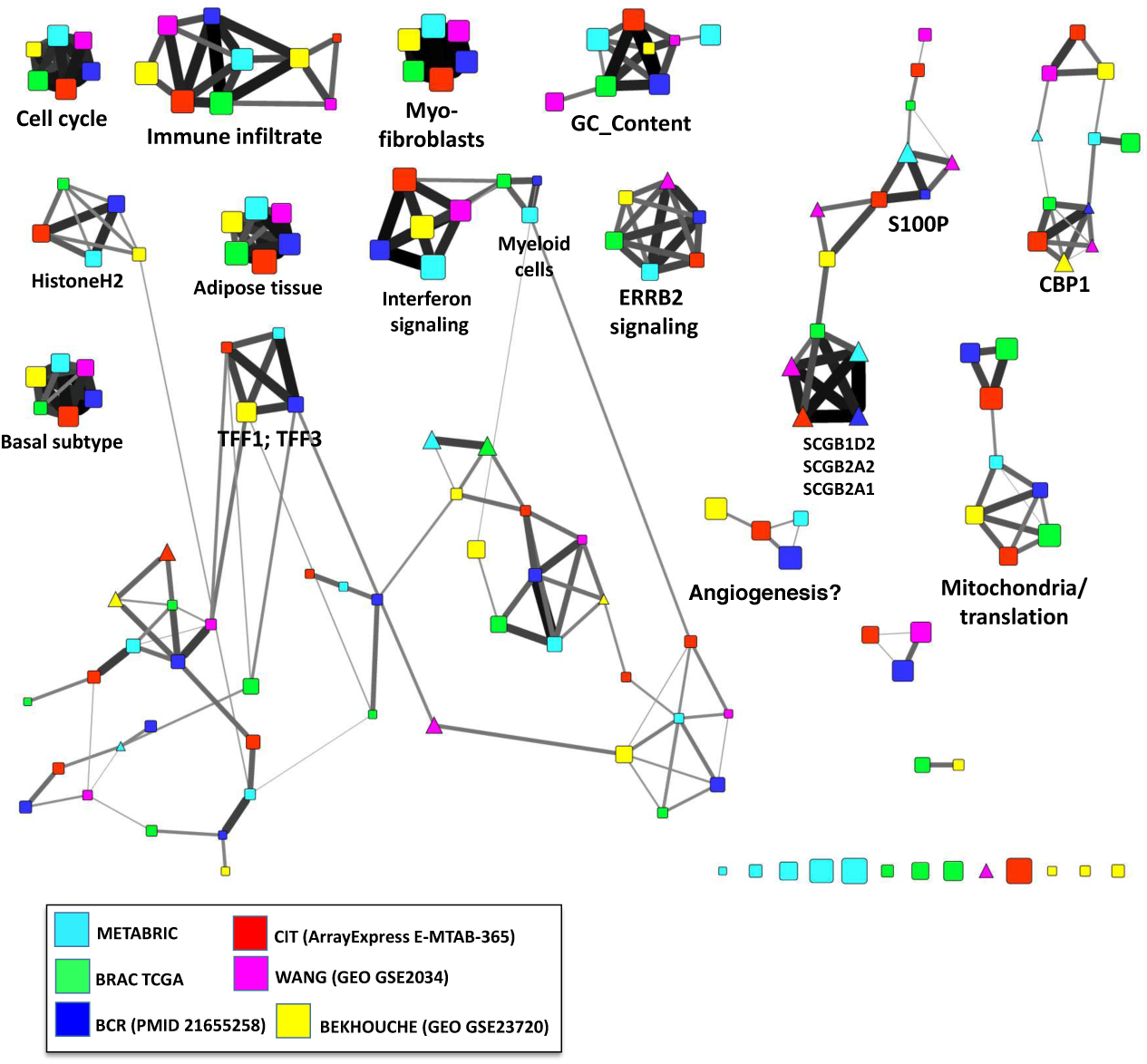
Graph of reciprocal correlations between components computed with MSTD choice for the reduced dimension and the number of components. The size of the points reflects their stability (larger points corresponds to more stable components). The color and the width of the edges reflect the Pearson correlation coefficient. Propositions of annotations of the pseudo-cliques in the graph are made based on the comparison with previously annotated metagenes (ref Biton) and the analysis of the top contributing genes using hypergeometric test and the *toppgene* web tool [28].

**Supplementary Figure SF4.**
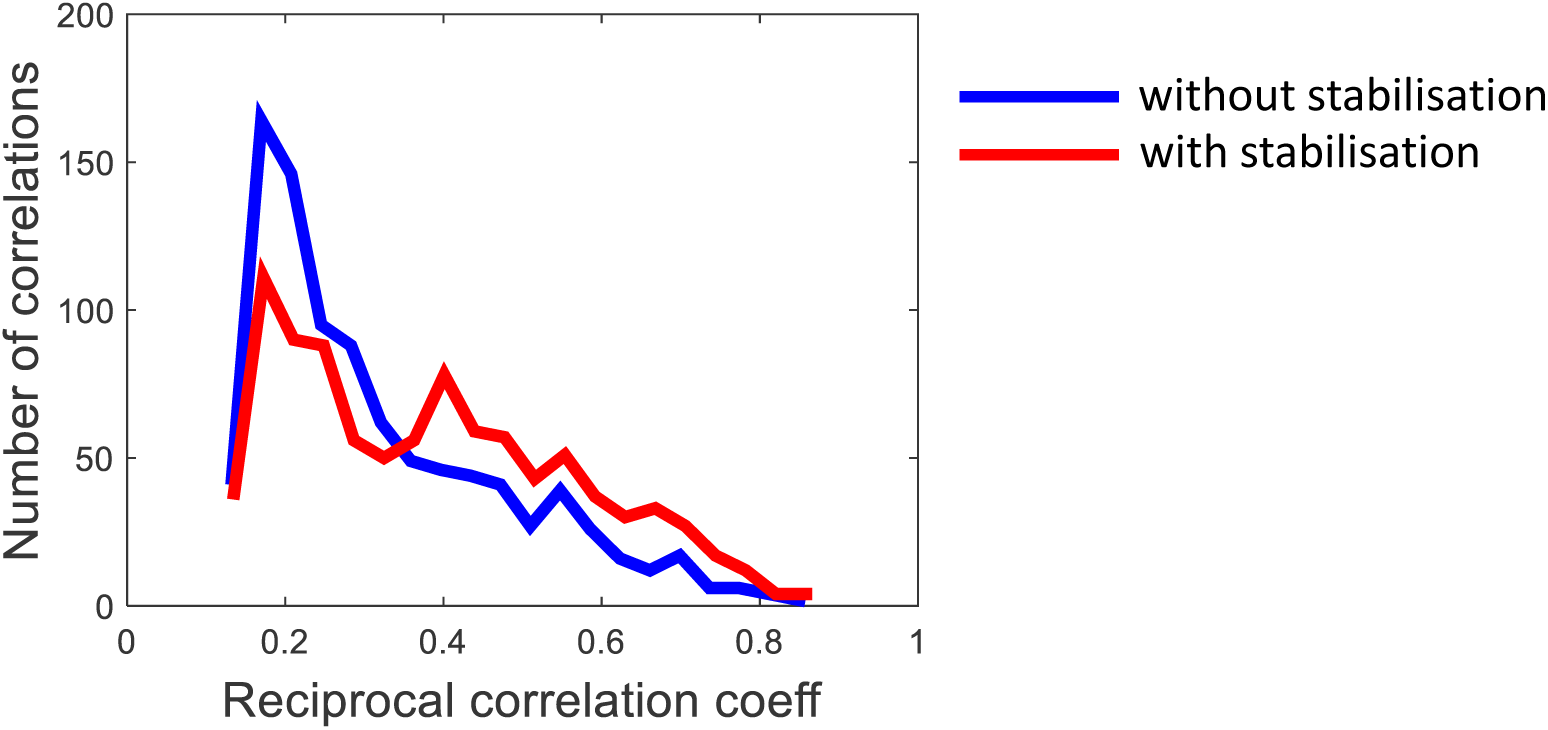
The histograms of the total number of reciprocal correlations in the correlation graph such as the one shown in Figure 3, with and without applying the component stabilization approach.

**Supplementary Figure SF5.**
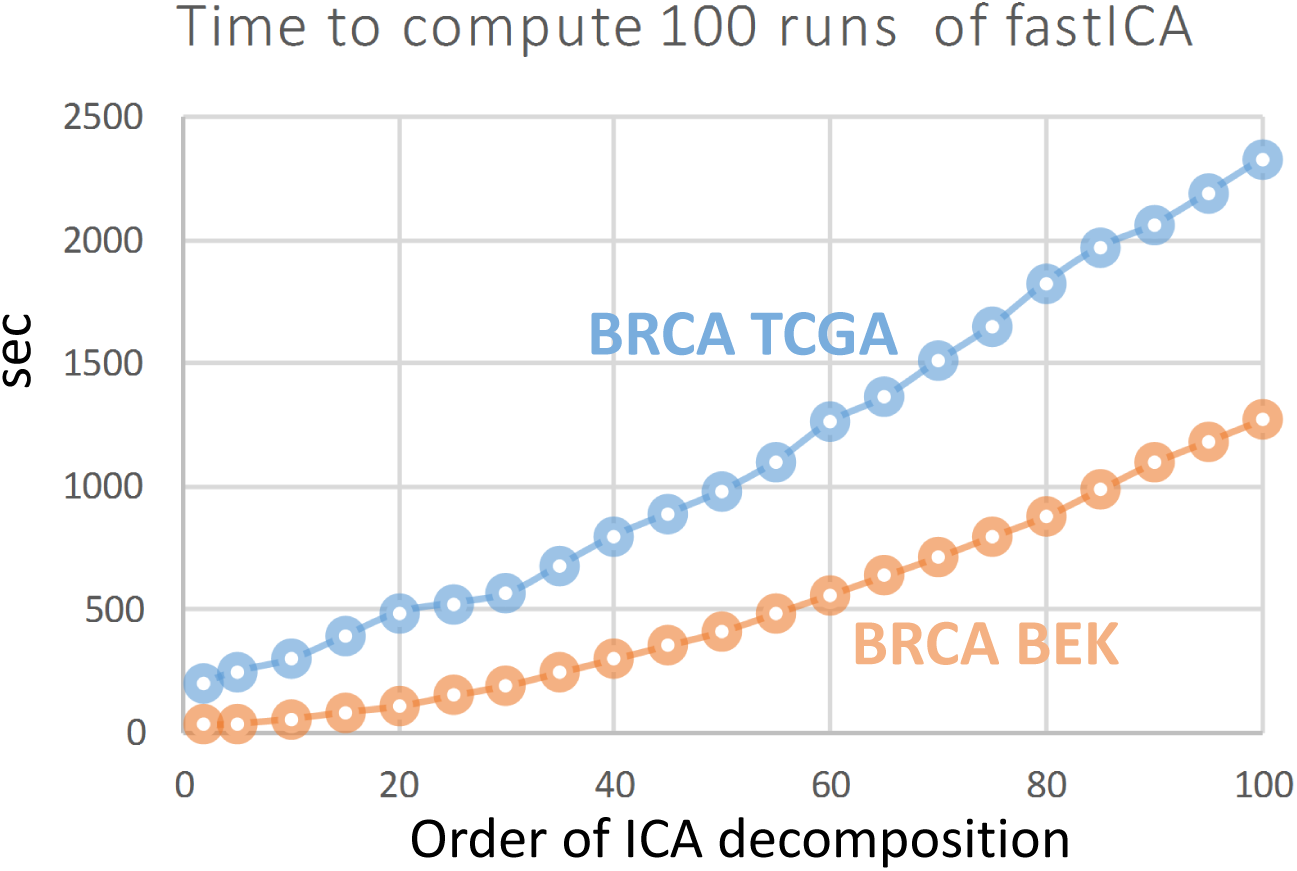
Computational time for ICA decomposition of different orders from 2 to 100 with step 5, using compiled MATLAB fastICA implementation and stability analysis by re-computing fastICA from 100 various initial conditions. The computation is made using an ordinary laptop with Intel Core i7 processor and 16Gb of memory, in a single thread. The BRCA BEK dataset (from [29]) contains 10000 genes in 197 samples, and the BRCA TCGA dataset (from [30]) contains 20503 genes in 1095 samples. The overall timing for computing all ICA decomposition with their stability analysis is 3.0 hours for BRCA BEK dataset, and 6.5 hours for BRCA TCGA dataset. These computations can be repeated from BIODICA software (https://github.com/LabBandSB/BIODICA), by launching ICA computation in scanning mode.

**Supplementary Table ST1.**
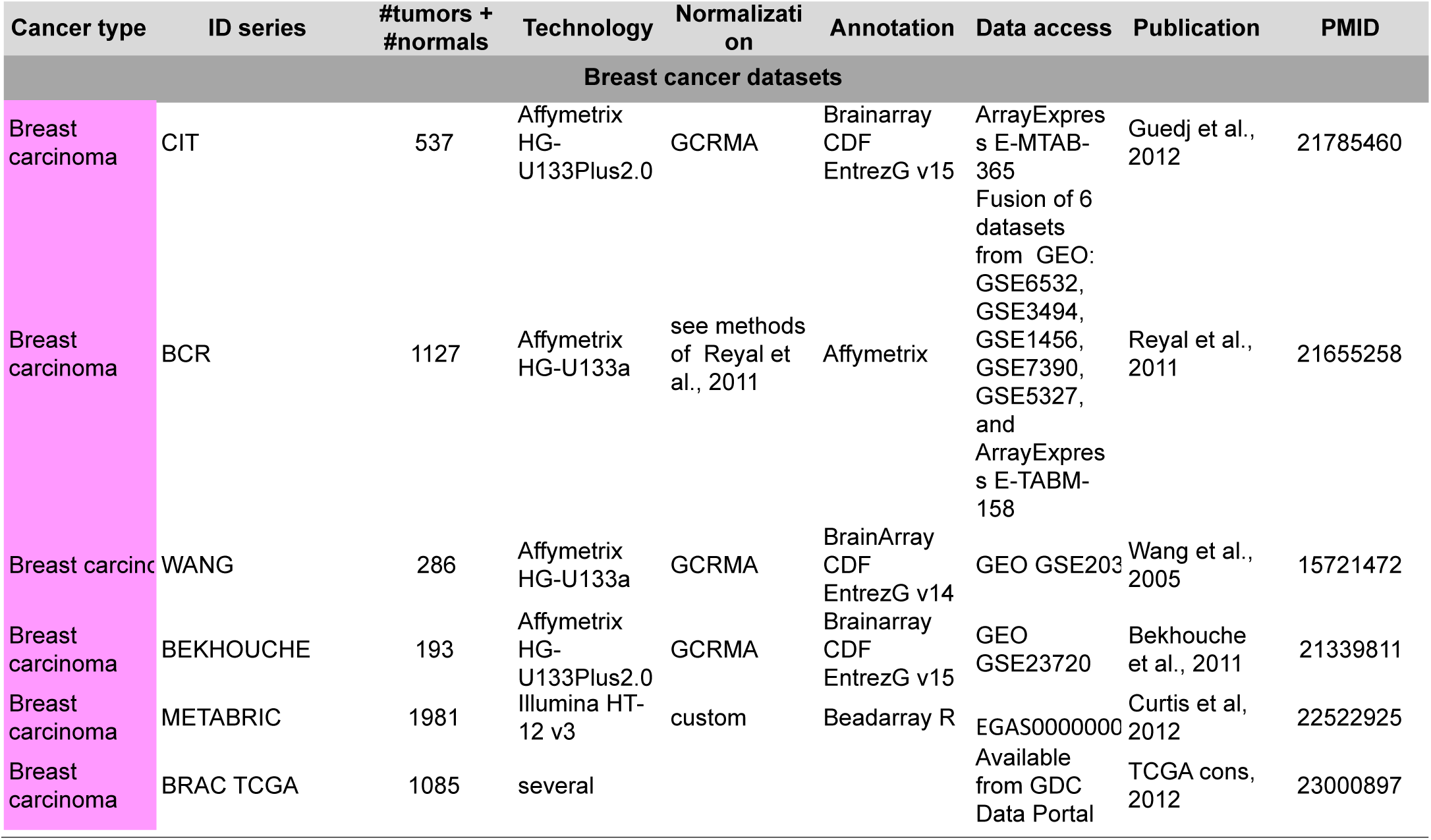
Breast cancer transcriptomic datasets used for the analysis of component reproducibility in independent datasets.

**Supplementary Table ST2.** Genes associated with ICA components of the METABRIC dataset, in the case when a component is driven by a small group of genes (frequently, one gene). Gene names marked in bold also drive independent components in several other breast cancer datasets and the corresponding components are reciprocally reproducible in terms of the correlation of the whole ICA-based metagenes.

|  |  |  |  |
| --- | --- | --- | --- |
| ABCC11 | **CPB1** | LTF | SCUBE2 |
| **ACOX2** | CST5 | MDK | SERHL2 |
| ADM | CXCL14 | MT1G | **SERPINA3** |
| AGR3 | CYP4X1 | MYBPC1 | **SERPINA5** |
| ALB | CYP4Z1 | **NAT1** | SHISA2 |
| ANKRD30A | DLK1 | NDP | SLC30A8 |
| **APOD** | DQ893812 | OLFM4 | SNAR-A3 |
| ASCL2 | EEF1A2 | PDZK1 | STC1 |
| ATP6V1B1 | GAGE12B | PEG3 | STC2 |
| AZGP1 | GAGE12I | PHGR1 | SYT13 |
| BC141517 | GAGE2B | PI15 | **TCN1** |
| C15orf48 | GAGE6 | PITX1 | TF |
| C19orf33 | GAGE8 | PLA2G16 | **TFF1** |
| C20orf114 | GRB14 | PRODH | **TFF3** |
| C4orf7 | HLA-A | **PSD3** | THRSP |
| C8orf84 | HLA-C | **PVALB** | UGT2B11 |
| CCND1 | HLA-DQB1 | RBP1 | UGT2B28 |
| CCNO | HLA-DRB1 | S100A7 | UGT2B7 |
| CD24 | HLA-DRB6 | S100A8 | VTCN1 |
| CENPV | HLA-H | S100A9 | WWOX |
| CFB | **HMGCS2** | **S100P** | X63359 |
| CKB | HOXB5 | **SCGB1D2** | XAGE1D |
| **CLEC3A** | LOC338579 | **SCGB2A1** |  |
| CLIC6 | LOC389493 | **SCGB2A2** |  |
| CNTNAP2 | LRRC26 | SCGB3A1 |  |

